# Seasonality of host-seeking *Ixodes ricinus* nymph abundance in relation to climate

**DOI:** 10.1101/2022.07.25.501416

**Authors:** Thierry Hoch, Aurélien Madouasse, Maude Jacquot, Phrutsamon Wongnak, Fréderic Beugnet, Laure Bournez, Jean-François Cosson, Frédéric Huard, Sara Moutailler, Olivier Plantard, Valérie Poux, Magalie René-Martellet, Muriel Vayssier-Taussat, Hélène Verheyden, Gwenaël Vourc’h, Karine Chalvet-Monfray, Albert Agoulon

**Author notes:** Current address: Ifremer, RBE-SGMM-LGPMM, La Tremblade, France.

## Abstract

There is growing concern about climate change and its impact on human health. Specifically, global warming could increase the probability of emerging infectious diseases, notably because of changes in the geographical and seasonal distributions of disease vectors such as mosquitoes and ticks. For example, the range of *Ixodes ricinus*, the most common and widespread tick species in Europe, is currently expanding northward and at higher altitudes. However, little is known about the seasonal variation in tick abundance in different climates. Seasonality of *I. ricinus* is often based on expert opinions while field surveys are usually limited in time. Our objective was to describe seasonal variations in *I. ricinus* abundance under different climates. To this end, a seven-year longitudinal study, with monthly collections of *I. ricinus* host-seeking nymphs, was carried out in France, in six locations corresponding to different climates. Tick data were log-transformed and grouped between years so as to obtain seasonal variations for a typical year. Daily average temperature was measured during the study period. Seasonal patterns of nymph abundance were established for the six different locations using linear harmonic regression. Model parameters were estimated separately for each location. Seasonal patterns appeared different depending on the climate considered. Western temperate sites showed an early spring peak, a summer minimum and a moderate autumn and winter abundance. More continental sites showed a later peak in spring, and a minimum in winter. The peak occurred in summer for the mountainous site, with an absence of ticks in winter. In all cases except the mountainous site, the timing of the spring peak could be related to the sum of degree days since the beginning of the year. Winter abundance was positively correlated to the corresponding temperature. Our results highlight clear patterns in the different sites corresponding to different climates, which allow further forecast of tick seasonality under changing climate conditions.

## Introduction

The effect of human activity on climate change is now recognised (IPCC, 2021), even though the extent of its impact remains open to debate. In particular, the influence of anthropogenic climate change on infectious disease incidence has to be assessed. Among infectious diseases, vector-borne diseases appear to be especially sensitive to climate change (Semenza and Suk, 2018) because changes in temperature and rainfall regime inevitably affect vector spatial distribution and phenology. However, climate change does not represent the unique disruptor of vector borne diseases (Rocklöv and Dubrow, 2020), which can also be impacted by biodiversity loss and land use change (Rizzoli et al., 2019).

Climate change may greatly affect the distribution of ticks, which are the main vectors of zoonotic pathogens in Europe (Gray et al., 2009; Ogden and Lindsay, 2016). Temperature influences tick physiology through the process of development, which is known to be accelerated with higher temperatures (Randolph et al., 2002). Tick host-seeking activity (referred to as “questing” for most tick species) also increases with temperature and relative humidity (Vail and Smith, 1998; Perret et al., 2003). Tick survival could, on the contrary, be reduced by higher temperatures and especially lower hygrometry (Daniel et al., 1976). The impact of climate change on tick distribution has already been observed and reported with a trend towards a northward expansion of *Ixodes ricinus* in Europe (Lindgren et al., 2000; Hvidsten et al., 2020) or its occurrence at higher altitude, in the Alps (Garcia-Vozmediano et al., 2020) or in mountain slopes in Norway (De Pelsmaeker et al., 2021).

Data on tick occurrence and climate variables have been confronted to define climate suitability for ticks such as *I. ricinus* (Estrada-Peña and Venzal, 2006). Extrapolation to future climate projections were then derived from these suitability maps (Estrada-Peña et al., 2012; Poretta et al., 2013). Besides these statistical models, mechanistic models incorporating knowledge on biological processes were also developed, for instance for *Hyalomma marginatum* (Estrada-Peña et al., 2011) or *I. ricinus* (Hoch et al., 2010). These process-based models could allow the simulation of the impact of climate change through the effect of meteorological variables on the involved processes. They represent relevant tools to predict the evolution of the transmission of tick-borne pathogens such as Crimean Congo Haemorrhagic Fever Virus (CCHFV) (Estrada-Peña et al., 2013; Hoch et al., 2018) or Lyme borreliosis agents, for which the northward expansion has been simulated in North America (Ogden et al., 2008). More recently, Li et al. (2016) developed a mechanistic agent-based model to simulate the seasonality of Lyme borreliosis risk, estimated through the number of infected nymphs in Scotland. Their model predicts an increase in disease risk at higher latitude and altitude, but also in the duration of *I. ricinus* host-seeking season, with warmer conditions. Such a longer duration of the host-seeking season has also been predicted by the model developed by Hancock et al. (2011), which focused on the evolution of the timing of *I. ricinus* peak of abundance in response to increasing temperature. Processes affecting the phenology of *Ixodes scapularis* were identified by a study linking simulation and data collection in different sites of the United States (Ogden et al., 2018). These authors identified different observed patterns of tick abundance depending on the site. They carried out several simulations considering different scenarios regarding temperature-independent diapause and development. The comparison of simulation outputs with observations suggested that diapause may be a major factor to explain geographical differences in tick phenology.

The objective of our study was to assess the differences in seasonal patterns of *I. ricinus* nymph abundance associated with distinct climates observed in France. A seven-year longitudinal study, involving monthly *I. ricinus* collections, was carried out at six locations in France. The species *I. ricinus* was targeted because it is responsible for major human vector-borne diseases in Europe (Lyme borreliosis and Tick Borne Encephalitis). This dataset allowed the fitting of statistical models of nymph abundance with time. Wongnak et al. (2022a) fitted a model on the same data set to explore the influence of meteorological variables on tick abundance for predictive purposes. Our modelling approach is complementary and aims to describe the seasonal patterns of *I. ricinus* by site and to relate them to climate features of the different geographical locations. Our objective was to assess these patterns for an average year, called typical year thereafter, by the use of the whole set of data for a given site, thus ignoring the between-year variations in the analyses. This approach can give insight into the evolution of tick phenology in relation to climate change.

## Data and models

### Tick sampling

Tick collection was carried out between April 2014 and June 2021 at six locations across France (Figure 1) which will be thereafter called Carquefou, Etiolles, Gardouch, La-Tour-de-Salvagny, Saint-Genes-Champanelle and Velaine-en-Haye. These locations were chosen in wooden habitats, selected for their high densities of host-seeking *I. ricinus* nymphs during spring, in different regions of France characterized by different climates (see Table S1, Supplementary Material, for GPS coordinates of the locations, altitudes and corresponding climates). In Gardouch, two nearby sites were sampled over the entire period: “Gardouch-in”, an enclosed forest patch with a fenced controlled population of roe deer, and “Gardouch-out”, another part of the same forest, just outside the fence of “Gardouch-in” and with uncontrolled roe deer populations. In La Tour-de-Salvagny, a first site (called “La Tour-de-Salvagny a” thereafter) was sampled between April 2014 and September 2016, then became inaccessible and was replaced by a second site (called “La Tour-de-Salvagny b” thereafter, approximately 2 km apart), sampled between April 2017 and June 2021. There was therefore a total of eight sites at six locations. Time-series of tick counts were obtained through 520 field campaigns carried out from April 2014 to June 2021, corresponding approximately to one collection session per month and per site (data were missing for 82 months due to several constraints: delay at the beginning, snow, Covid-19 lockdown…). At each site, ten marked transects of 10 x 1 m long were chosen along trails covered by short grass or a leaf litter, with a minimum distance of 20 m between them. Each tick collection session corresponded to a single day, selected for the absence of rain or snow and the dryness of the grass and leaf litter cover. On that day, on each transect, a 1 m^2^ white flannel was slowly dragged along the 10 m to collect host-seeking ticks (Agoulon et al., 2012): ticks were counted, removed from the cloth with tweezers and stored alive for further identification at stage and species level according to identification keys (Pérez-Eid, 2007). Tick collection was repeated three times consecutively on the same marked transects in order to collect more individuals and to improve accuracy in the assessment of tick abundance (Bord et al., 2014). This method simulates the natural detection and infestation of a host by host-seeking ticks. The number of collected ticks was therefore considered representative of the host-seeking tick abundance on the day of collection. Due to a low number of adult ticks collected and the known low involvement of larvae in disease transmission cycle, these development stages were not considered for further analyses: only *I. ricinus* nymphs were considered, reflecting the major risk of pathogen transmission to humans (Kurtenbach et al., 2006).

**Figure 1:**
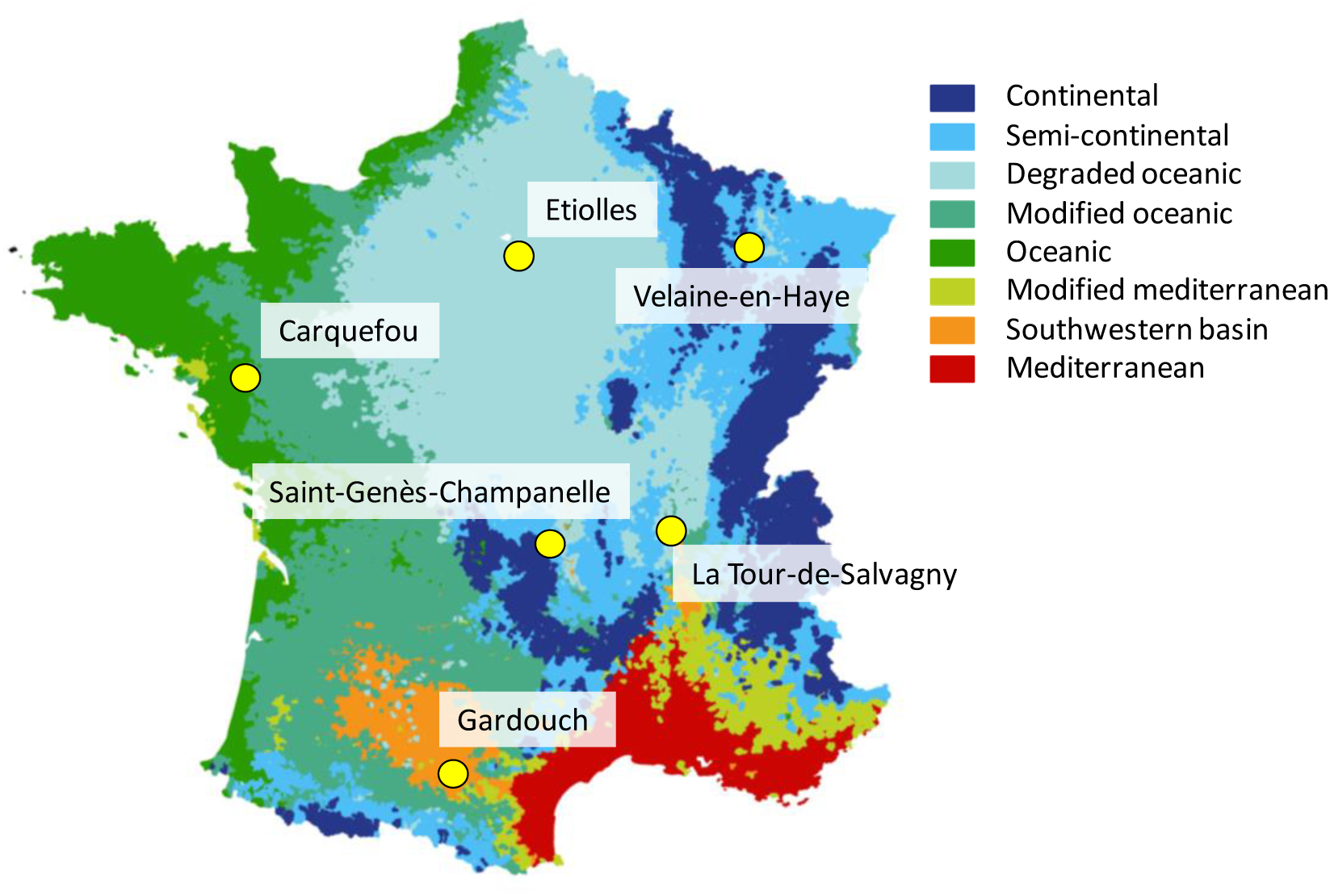
Map of France with the different climates encountered (from Joly et al., 2010). Tick sampling locations are indicated

### Local meteorological data

Weather stations were installed at the beginning of the period at each location to collect hourly air temperature and relative humidity records at a height of 1.5 meters. A single station was used for La Tour-de-Salvagny a and b, and for Gardouch-in and Gardouch-out. Daily mean meteorological variables were obtained from those records. When data were missing, data from nearby MeteoFrance and INRAE automatic weather stations were collected to impute the missing values, with a random forest approach (Wongnak et al, 2022).

Daily Vapor Pressure Deficit (VPD, in kPa) was computed by the following equation:

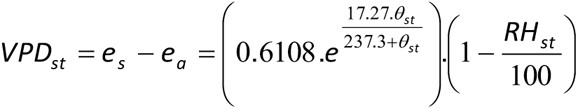

where *s* and *t* indices represent the respective site and recording date, *e_a_* is the amount of moisture in the air, *e_s_* the maximum water holding capacity, *θ* the temperature and *RH* the relative humidity.

For each location, the daily mean temperatures and VPDs of an average year were obtained by averaging the corresponding daily mean meteorological data from 1 January to 31 December (see Figure S1 and S2, Supplementary Material).

In order to assess the influence of temperature on the date of the estimated peak, a cumulative temperature (in degree-day) was computed for each location. It was calculated by summing the daily mean temperatures of an average year from the 1^st^ January up to the estimated date of the peak.

### Statistical analyses

The objective of the analyses was to describe the site-specific seasonal variations in the abundance of *I. ricinus* nymphs, hereafter referred to as tick abundance pattern. The distributions of the numbers of nymphs for each site were right skewed, typically with a mode below 10 nymphs collected per day and maximum values that could reach several hundred. Taking the natural logarithm of the number of nymphs allowed to make the distributions more symmetrical (see Figure S3, Supplementary Material). The abundance patterns were modelled using harmonic regressions. Briefly, harmonic regression is a type of linear regression in which the days of the year are mapped onto a circle, and, functions of the sine and cosine of the corresponding angles are included as covariates in the regression.

The outcome of all models was the natural logarithm of the number of *I. ricinus* nymphs (+1 to avoid ln(0), which is not defined) collected at a given site on a given day. This outcome was modelled as a function of day of the year at the time of collection. The models’ specifications were as follows:

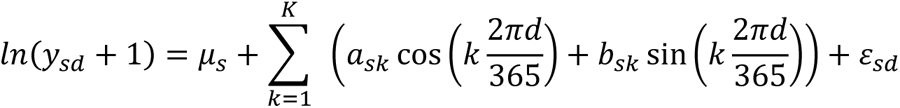

with:

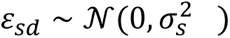

Where *y*_*sd*_ was the number of nymphs collected at site *s* on day of year *d* (ranging from 1 to 365); μ_*s*_was the model intercept for site *s*, *K* was the maximum number of periodic (also called Fourier) terms included; and ε_*sd*_ was the residual error with mean 0 and variance σ^2^.

Increasing *K* makes it possible to model increasingly complex seasonal patterns. In order to mitigate the risk of overfitting with high values of *K*, models with *K* varying between 0 and 4 were compared based on the Akaike Information Criterion (AIC) value. For most locations, there was an important decrease in AIC between *K* = 0 and *K* = 2 (see Figure S4, Supplementary Material) and we reach a plateau thereafter. Therefore, *K* = 2 was used in all models.

Using the fitted models, typical within-year evolutions of tick abundance were simulated. The characteristics of interest included the date of peak abundance, i.e. the date when the predicted number of *I. ricinus* was highest. The relation of the peak dates with cumulative mean temperatures were investigated.

Unlike peak tick abundance, which depends on many factors which have not been accounted for in the model (e.g. host abundance), we hypothesize that winter tick abundance relates to corresponding temperature, which has an influence on tick activity. To analyse these potential links, the average January daily mean temperature was related to the corresponding estimated mean tick abundance. To improve comparability among sites, average tick abundance in January was standardized by its maximum value in each site. Similarly, as VPD is recognized to lower tick activity (Vail and Smith, 1998), we linked the average August VDP to the corresponding relative tick abundance.

## Results

### Seasonal patterns of observed I. ricinus nymph abundance at the different sites

Figure 2 shows the natural logarithm of the number of *I. ricinus* nymphs (+1) collected as a function of the day of the year, for the eight sites and seven years of collection. At a given site, seasonal patterns were stable over years, whereas there were variations between sites exhibiting different patterns. Therefore, we fitted different models for the different locations, including data from all years for each site. When there were two sites for the same location (*i.e.* Gardouch in and out, La Tour-de-Salvagny a and b), the patterns looked similar. In those cases, we included site as a fixed effect in the harmonic regression (different mean for each site), but the same coefficients were used for the Fourier terms: this means that the fitted curves followed the same seasonal dynamics but were of different heights.

**Figure 2:**
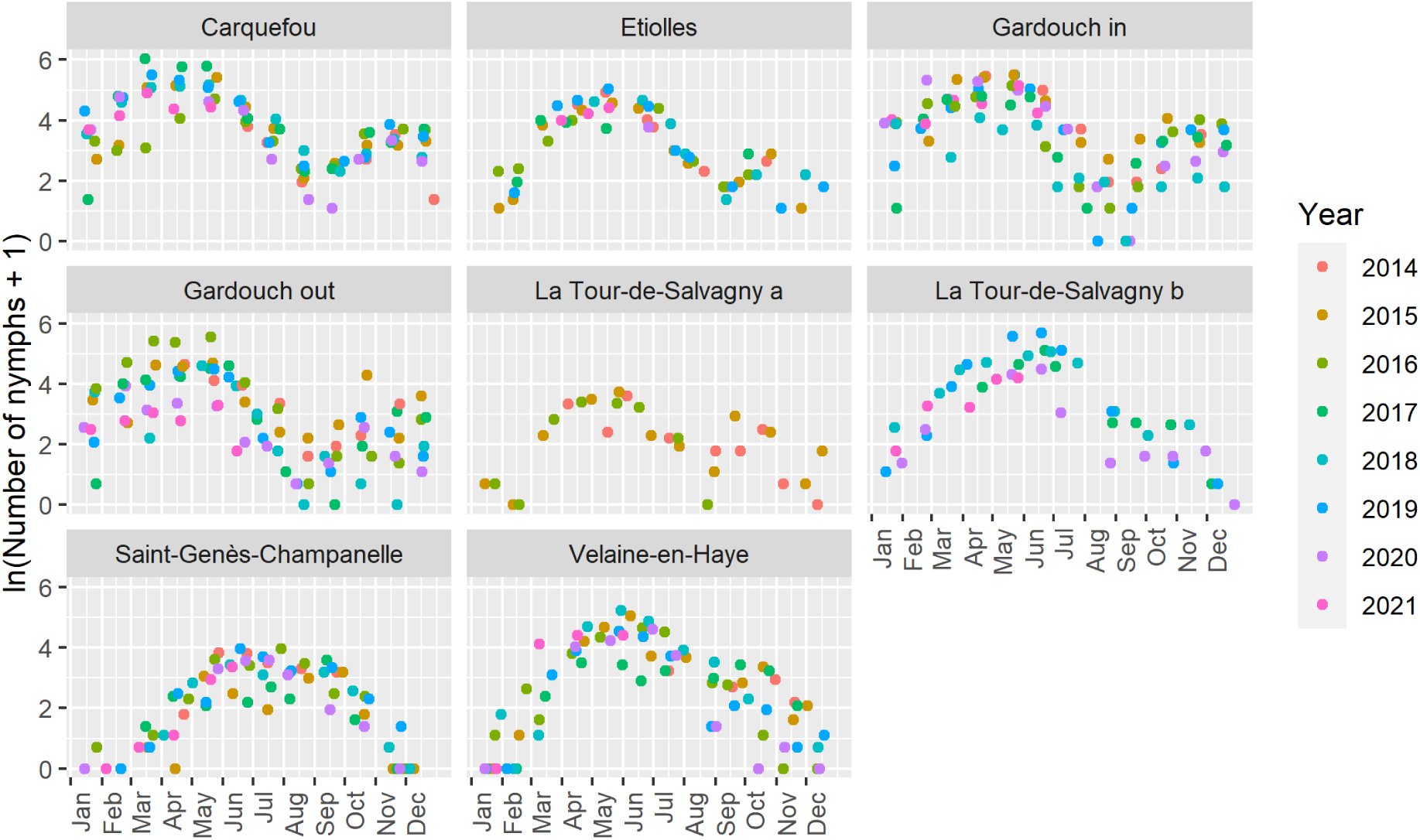
Natural logarithm of the observed number of *Ixodes ricinus* nymphs (+1) collected as a function of day of the year, at eight French sites over seven years.

### Seasonal patterns of fitted I. ricinus nymph abundance at the different sites

The different models appear to correctly represent the observed dynamics (Figure 3) and particularly the spring peak. For Gardouch-in and Gardouch-out as well as for the two sites of La Tour-de-Salvagny, the choice of fitting the same model with a distinct intercept seems relevant. In particular, for the latter, the model succeeds in representing both the same dynamics and the difference in abundance between the two sites. When applied over the whole data set without accounting for the sampling year, such a modelling helps to identify the favourable and less favourable years for tick abundance, for which observed tick abundances are respectively above and below the fitted curve. Interestingly, favourable years seem to differ from one site to another, even on the same location (Gardouch in and out). For instance, the model revealed a favourable year in 2016 at Gardouch out (but not at Gardouch in), in 2017 at Carquefou, in 2018 at Velaine-en-Haye and in 2019 at La Tour-de-Salvagny (b).

**Figure 3:**
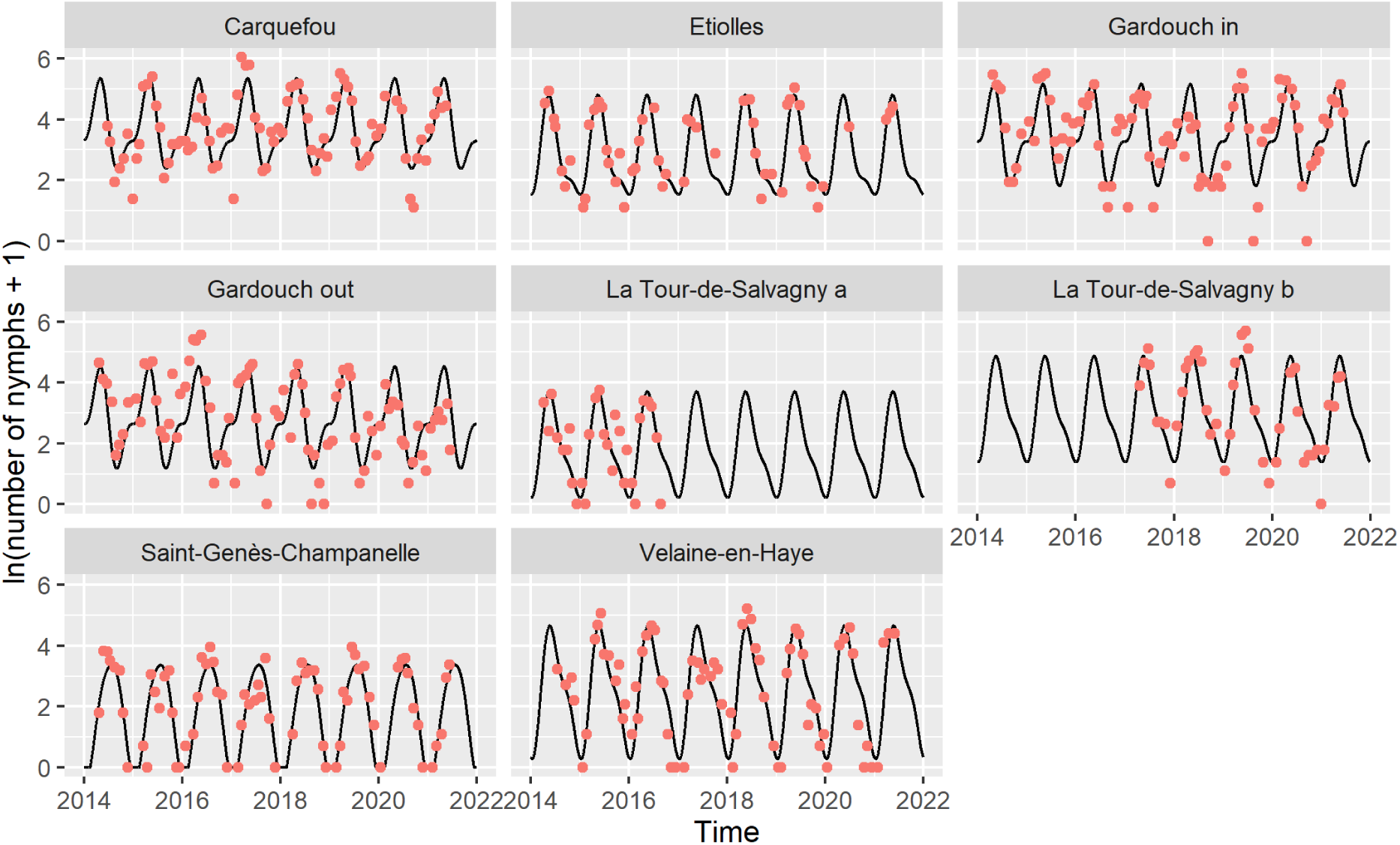
Natural logarithm of the number of *Ixodes ricinus* nymphs (+1) predicted by the models (black curves) and number of nymphs effectively collected (red dots), at eight French sites over seven years.

To compare sites, we superimposed the predicted tick abundance for all sites over one year (Figure 4). The predicted patterns show a spring peak in all sites except Saint-Genès-Champanelle, where the maximal abundance is observed in summer. Some small differences in spring peak dates are predicted for the other sites. Tick peak occurs earlier for the western “oceanic” sites (Carquefou and Gardouch) and later for the eastern “continental” sites (Etiolles, La Tour-de-Salvagny and then Velaine-en-Haye). Winter abundance highlights another striking feature: in the western sites (Carquefou and Gardouch), predicted tick abundance is increasing from September to November to reach a plateau throughout the winter months, which is not observed elsewhere.

**Figure 4:**
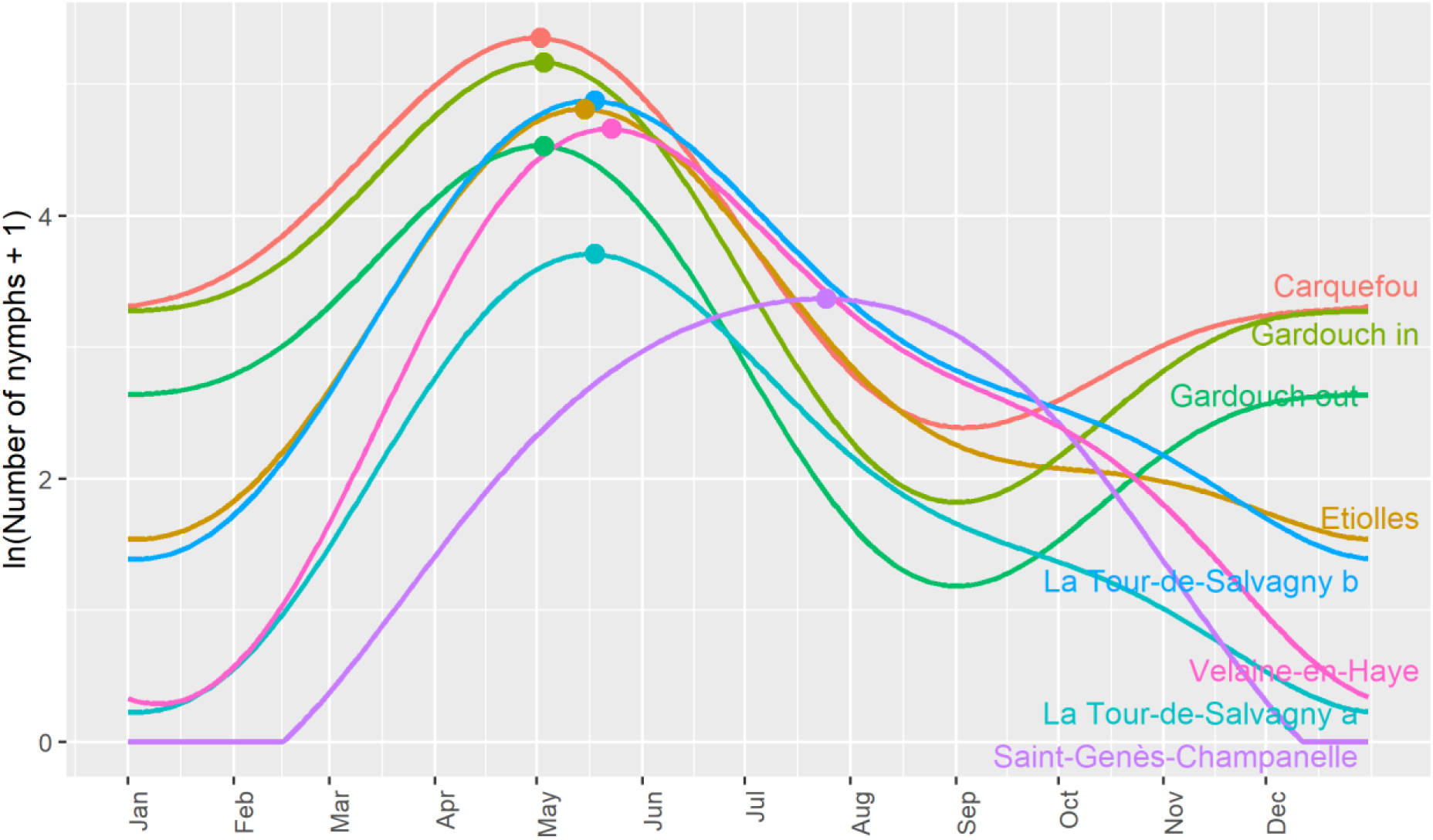
Predicted annual log-transformed abundance patterns of host-seeking *Ixodes ricinus* nymphs for the different sampling sites. The dots represent the maximum of each curve, i.e. the peak of abundance.

### Links between predicted tick abundance and meteorological variables

The predicted spring peak date can be related to the cumulative sum of temperatures from the beginning of the year, hence resulting in a degree-day effect (Figure 5). The order by which sites are ranked with respect to the cumulative temperature corresponds to the order of spring peak dates. The simulation exhibits an early peak in Gardouch and Carquefou, followed by Etiolles, La Tour-de-Salvagny, Velaine-en-Haye and Saint-Genès-Champanelle. For the four sites with the earliest spring peak, the peak occurs when temperature cumulative sum reaches c.a. 1000 °C.d (Table I). At the more continental site of Velaine-en-Haye, the maximum is reached at a lower value for the cumulative temperature. The mountainous site, Saint-Genès-Champanelle, appears to have a different behaviour regarding temperature, with a cumulative temperature around 1600 °C.d at peak date.

**Figure 5:**
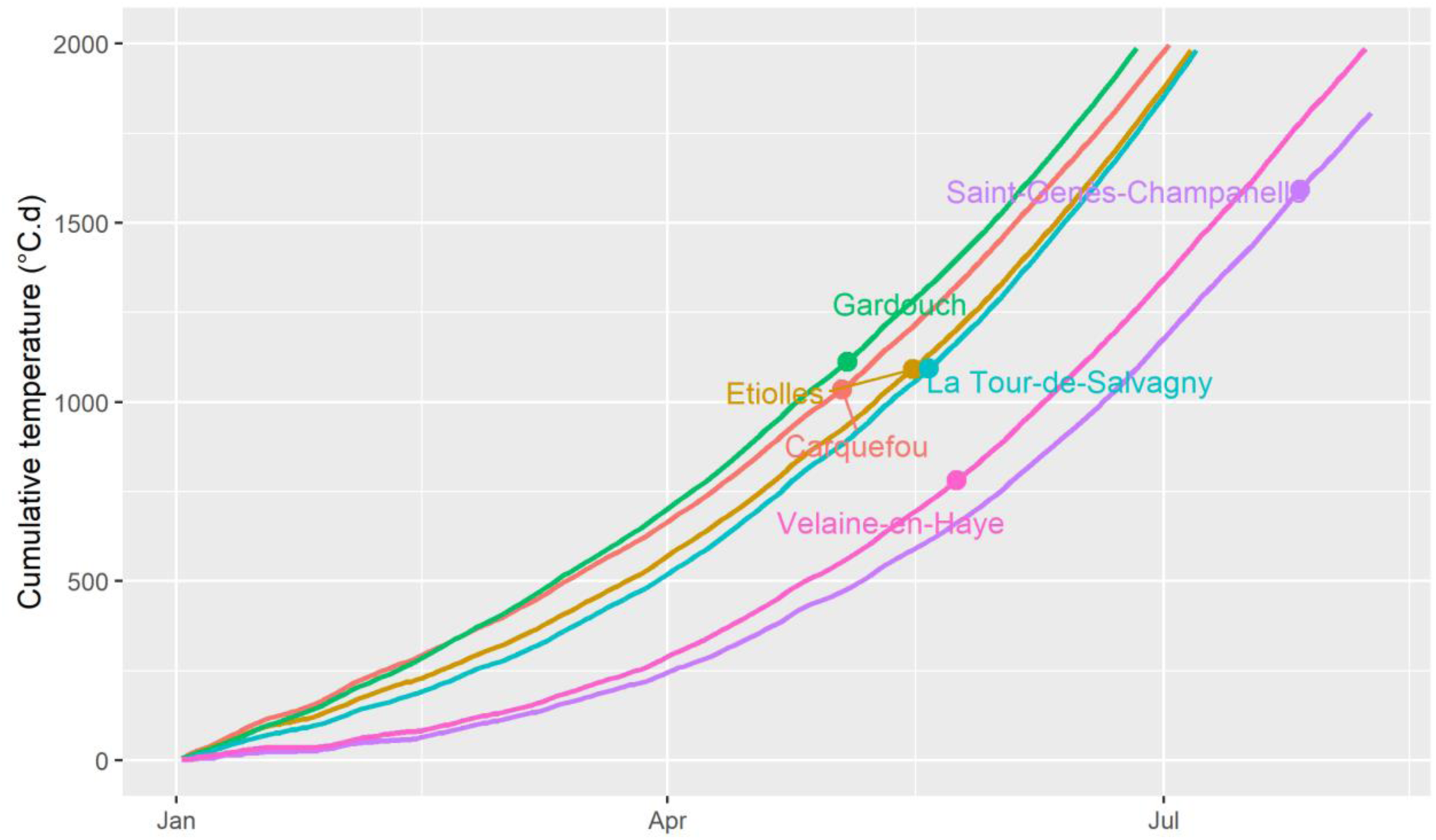
Cumulative temperatures over time for the six locations, with dots corresponding on the X-axis to the predicted dates of *I. ricinus* host-seeking nymph peak abundance at each location.

**Table I:** Peak date (predicted date of *I. ricinus* host-seeking nymph peak abundance), cumulative tem perature (°C.d) and mean daily temperature (°C) at peak date for the different locations.

|  | Carquefou | Gardouch | Velaine-en-Haye | La Tour-de-Salvagny | Etiolles | Saint-Genès-Champanelle |
| --- | --- | --- | --- | --- | --- | --- |
| Peak date (d) | May 2<br>(122) | May 3<br>(123) | May 23<br>(143) | May 18<br>(138) | May 15<br>(135) | July 25 (206) |
| Cumulative temperature (°C.d) at peak date | 1034.5 | 1112.8 | 781.5 | 1093.3 | 1092.1 | 1593.2 |
| Mean daily temperature (°C) at peak date | 11.5 | 12.6 | 12.7 | 14.1 | 12.5 | 17.2 |
| Average January nymph abundance | 0.637 | 0.643 (in)<br>0.593 (out) | 0.076 | 0.086 (a)<br>0.305 (b) | 0.338 | 0. |
| Average January daily temperature (°C) | 6.55 | 6.04 | 1.65 | 4.06 | 5.03 | 1.37 |

Table I shows the range of values of the mean daily temperature at the time of the peak between the different sites. The mountainous site of Saint-Genès-Champanelle shows again a clear different behaviour regarding temperature, with a peak of tick abundance corresponding to summer temperature (17.2°C). For the other sites, the peak of abundance is obtained in spring for intermediate temperature values, *i.e.* between 11.5°C (in Carquefou) and 14.1°C (in La Tour-de-Salvagny).

Moreover, average predicted tick abundance in January, standardized by the yearly maximum value at the corresponding site, can be directly related to the average temperature for the same period (figure 6). This straightforward significant relationship shows that mild winter temperatures of Carquefou and Gardouch correspond to higher winter tick abundance than other sites.

**Figure 6:**
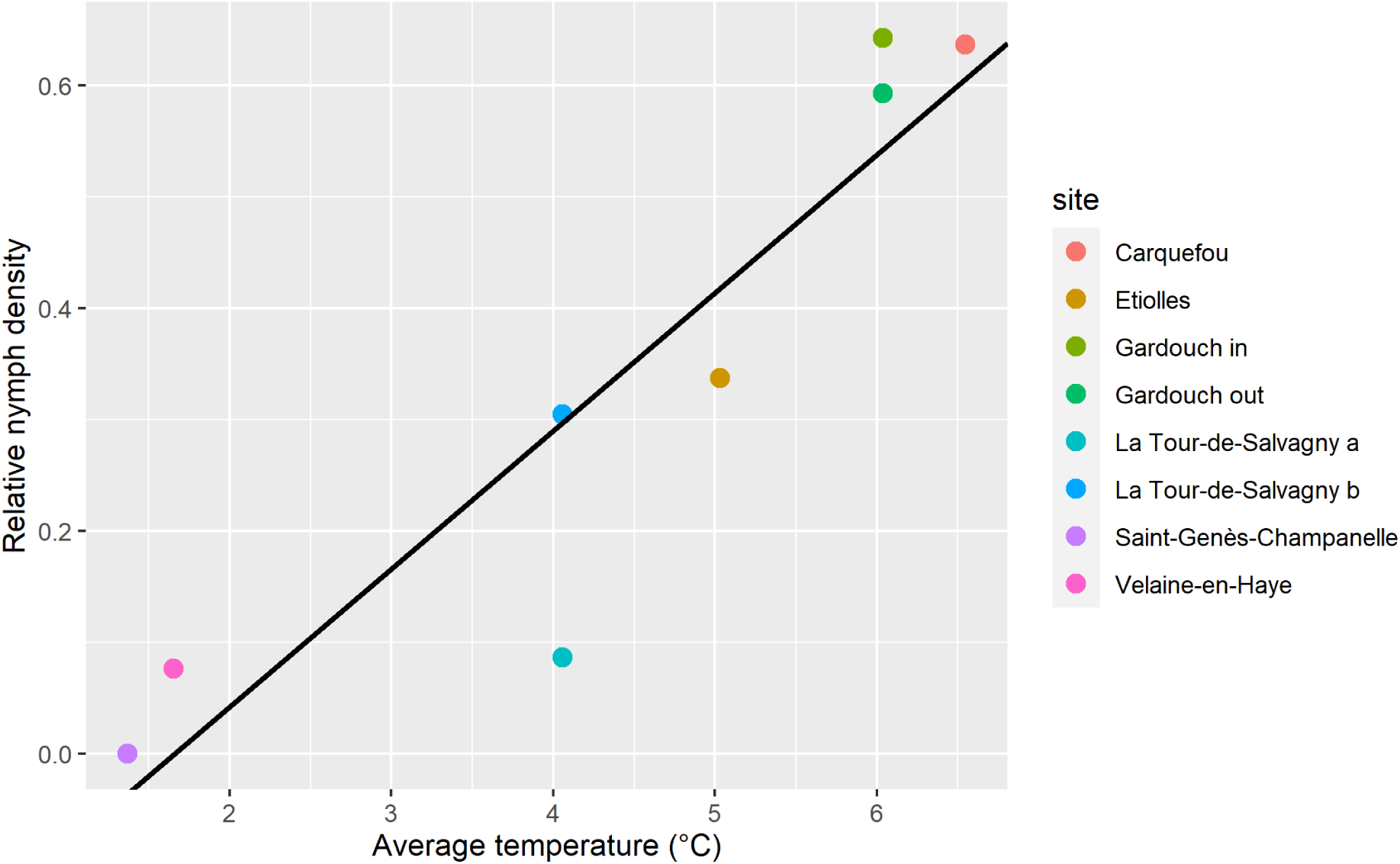
Predicted mean nymph abundance in January, standardized by the yearly maximum value, as a function of corresponding mean temperature in January for the different sites. Equation of the regression line: *y* = −0.206 + 0.124*x*, p=0.001.

On the contrary, only a weak relationship was shown between average daily mean VPD and relative tick density (see Figure S5, Supplementary Material), with an estimated Pearson coefficient of –0.24.

## Discussion

The tick abundance patterns modeled in this work can be divided into three types: two types exhibiting a unimodal curve, with a peak either in spring (Etiolles, La Tour-de-Salvagny, Velaine-en-Haye) or in summer (mountainous site of Saint-Genès-Champanelle), and a third type exhibiting a bimodal pattern in Carquefou and Gardouch, with a peak in spring, a minimum in summer and an increase with a plateau in autumn and winter. When comparing sites that exhibit a spring peak, the predicted peak occurs later as the site is located further east. Collected tick abundance declines thereafter in summer in these sites, which might be due to a decrease in the host-seeking nymph population (due to the fixation of nymphs on hosts in the spring) or a decrease in the questing behaviour of the remaining nymphs due to a lower relative humidity in summer. These fitted tick abundance patterns are consistent with the literature. We used the same data set as Wongnak et al. (2022), who derived patterns applied to the different sites. However, their results relied on a descriptive analysis of the data, whereas our conclusions are based on a statistical model using harmonic regression relating tick abundance with time, thus dedicated to the assessment of tick phenology. Even though both analyses often converge, some differences may be highlighted, as for instance the absence of any autumn peak in eastern sites in our analysis. Several field studies highlight differences in tick phenology according to climate. Thanks to seven year monthly samplings, Alonso-Carné et al. (2015) depicted a unimodal or a bimodal curve in two locations in Spain differing from their respective climatic characteristics (continental *vs* temperate). From Bregnard et al. (2021a) tick phenology is expected to be unimodal at northern latitudes and bimodal in areas with warmer and longer summers. They also reported 12 different studies on tick phenology across Europe where the authors identified a unimodal or a bimodal phenology, but observed phenology does not always follow these rules, possibly due to year-to-year variations. Babenko (1958) (in Korenberg, 2000) depicted four stylized types of seasonal changes in the abundance of *I. ricinus* adults. More specifically, he considered bimodal curves for temperate zones (*i.e*. Crimea) and unimodal ones with a peak in summer for continental areas of Russia. Even applied to adult ticks, these curves can be compared with our predictions for the nymph stage in Carquefou and Gardouch (oceanic and southwestern basin climate respectively), and in Saint-Genès-Champanelle (continental climate), respectively. Concerning the influence of altitude, differences in tick phenology were shown between transects situated respectively at 100 and 400 meter high in Norway (Qviler et al., 2014). In Switzerland, Bregnard et al. (2021a) showed a cline ranging from bimodal to unimodal curves when reaching higher altitude (altitude ranging from 620 to 1073 m in their study). According to Gray et al. (2016), *I. ricinus* abundance patterns for larvae, nymphs and adults follow a bimodal curve in Central and Western Europe, but these authors also point out that there are variations on this expert-based pattern. Our fitted model based on harmonic regression and using longitudinal data resulting from 8 years of monthly surveys allows us to relate these seasonal variations to observations. It also suggests that these two types of patterns are both present in different locations of the French territory.

The abundance patterns can be directly related to what we know about the specific climate of the different sites. A gradient in spring peak date is exhibited from the western sites located in temperate areas to the more eastern ones, corresponding to a semi-continental climate (see the climate typology in France from Joly et al., 2010 on figure 1). Mountainous areas can be assimilated to even more continental zones, with however a heat deficit during summer. An early spring peak corresponds to temperate sites where temperatures are warmer in winter and at the beginning of spring. Diuk-Wasser et al. (2006) also highlighted an earlier peak for *Ixodes scapularis* nymphs in southern vs northern sites in USA. Temperature has long been identified as the major factor acting on tick development (Randolph, 2004), each stage necessitating a certain amount of degree-days to develop to the next stage (Hoch et al., 2010). The spring peaks in the different sites logically occur successively in accordance with the cumulative temperature curves (figure 5). Four locations (Carquefou, Etiolles, Gardouch and La Tour-de-Salvagny) exhibit a spring peak for comparable degree-day values (around 1000 °C.d). This is not the case in Saint-Genès-Champanelle (around 1600 °C.d) and to a lesser extent in Velaine-en-Haye (around 800 °C.d). For the mountainous location, in Saint-Genès-Champanelle, a peak is hardly identified in summer and the tick density curve looks smoothed, which generates uncertainty in the estimation of the peak date: the amount of degree-days may be over-estimated. Temperature data measured at 1.5 meter from the ground may not represent the temperature experienced by the ticks at ground level. Furthermore, factors other than cumulative temperature may influence the date of the peak. The date of the peak may depend on the proportion of larvae which are developing into nymphs during the spring: if development from larvae to nymphs occurs before the spring, *i.e.* in the autumn of the previous year (as suggested by Randolph et al., 2002), resulting host-seeking nymphs may be active very early in the spring, at the onset of warming. It has also been shown that the host-seeking behaviour (questing) of different tick populations may respond differently to temperature, depending on the considered climate (Tomkins et al., 2014). Ticks that have achieved their development start questing with a rate that depends on relative humidity and temperature. According to Beugnet et al. (2009), this proportion of questing ticks is maximal for intermediate temperatures, *i.e.* between 10 and 20°C. A field study also reported maximum values of collected ticks for intermediate temperatures in Germany (Gethmann et al., 2020), *i.e.* 13-15°C of ground temperature, which is comparable to our mean daily air temperature at peak date (table I). Our values for temperature at the peak are consistent with these expert– and data-based questing rates. Finally, a clear relationship has been shown between winter (January) abundance in host-seeking nymphs and corresponding average temperature in the different sites. This is supported by findings of Furness and Furness (2018), who showed a winter increase in the infestation of birds by immature *I. ricinus* ticks with temperature. As stated by Gray et al. (2009), climate change is likely to increase the duration of the host-seeking period of ticks, thus highlighting the need of winter tick collections to monitor the risk of tick bite.

A weak relationship was put in evidence between August tick abundance and VPD. Perret et al. (2000) found a higher negative relationship between the evolution of monthly nymph abundance between April and September and the corresponding variation in VPD. Although the relationship could be due to a negative effect of high VPD on tick activity, which has been experimentally demonstrated by Vail and Smith (1998) with a weekly sampling, it also results from a natural decline of questing tick abundance after the spring peak. In our study, a between-site effect of VPD could possibly not be shown due to our monthly sampling since tick host-seeking activity in summer may be highly fluctuating due to very variable conditions of hygrometry, especially at ground level.

Concerning inter-annual variations in tick abundance, we note that years with relatively high observed abundance compared with simulated abundance are not the same for different sites. While these variations can be explained by inter-annual variations of local meteorological conditions, other factors may also be involved, including host densities. Nymph abundance is known to be related to the rodent abundance of the previous year, as shown by Perez et al. (2016) for the effect of wood mice (*Apodemus sylvaticus*) abundance on *I. ricinus* densities. The indirect effect of beech fructification two years before on nymph density and acarological risk (density of infected nymphs) has also been demonstrated respectively by Brugger et al. (2018) and Bregnard et al. (2021b).

In this study, tick abundance at different sites in France showed repeatable within year patterns across an 8-year period. At a given site, inter-annual variation was usually low compared with seasonal variation. The identified associations between climate characteristics and seasonal patterns of tick abundance corroborate expert opinions. Peak abundance usually occurred for cumulative temperatures of around 1 000°C, with extremes of 780°C and 1600°C. The different profiles identified at the scale of France are a first step towards the understanding of tick phenology and its variation according to climate, and give new insight into the influence of climate change on tick activity that has to be put in perspective with disease risks.

## Data availability

The datasets generated during and/or analysed during the current study are available in the “TEMPO— Réseau National d’Observatoires de la Phénologie” repository, https://doi.org/10.15454/ZSYGUM.

## Acknowledgements

Tick observatories were supported by the CC-EID and Climatick projects: “Adaptation of Agriculture and Forests to Climate Change” metaprogramme of the French National Research Institute for Agriculture, Food and Environment (INRAE). We also would like to thank the TEMPO French Network (https://tempo.pheno.fr/) for providing resources to support this study.

