## Supplementary Material for "Seasonality of host-seeking *Ixodes ricinus* nymph abundance in relation to climate"

**Supplementary Informations**

| Location | Position (latitude, longitude) | | Altitude (m) | Climate (Joly et al., 2010) |
|---|---|---|---|---|
| Carquefou | 47.319461 | -1.491685 | 7.4 | Oceanic |
| Etiolles | 48.667431 | 2.479903 | 84.2 | Degraded oceanic |
| Gardouch | 43.370952 | 1.673424 | 238.3 | Southwestern basin |
| La Tour-de-Salvagny | 45.797595 | 4.712050 | 286.2 | Degraded oceanic |
| Saint-Genès-Champanelle | 45.705787 | 2.971739 | 966.8 | Continental |
| Velaine-en-Haye | 48.704871 | 6.082540 | 334.1 | Semi-continental |

Table S1: Position, altitude and corresponding climate of the different sampling locations

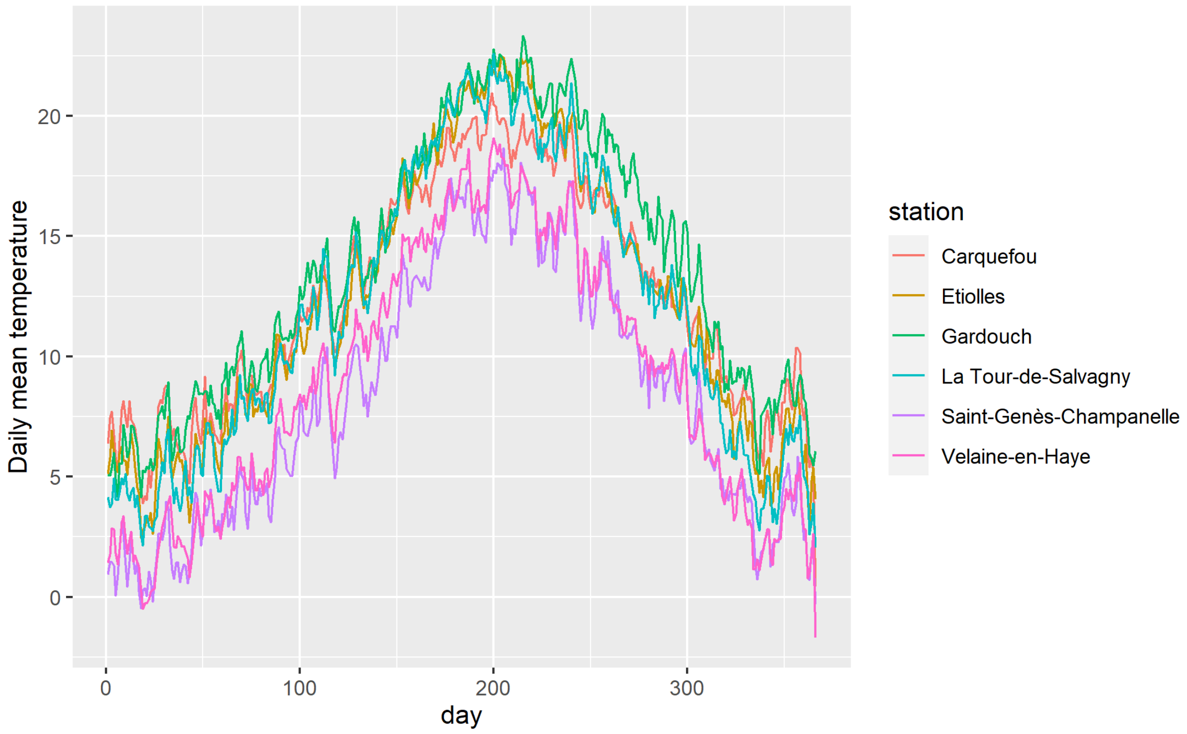


Figure S1: Daily mean observed temperature data for the different locations.

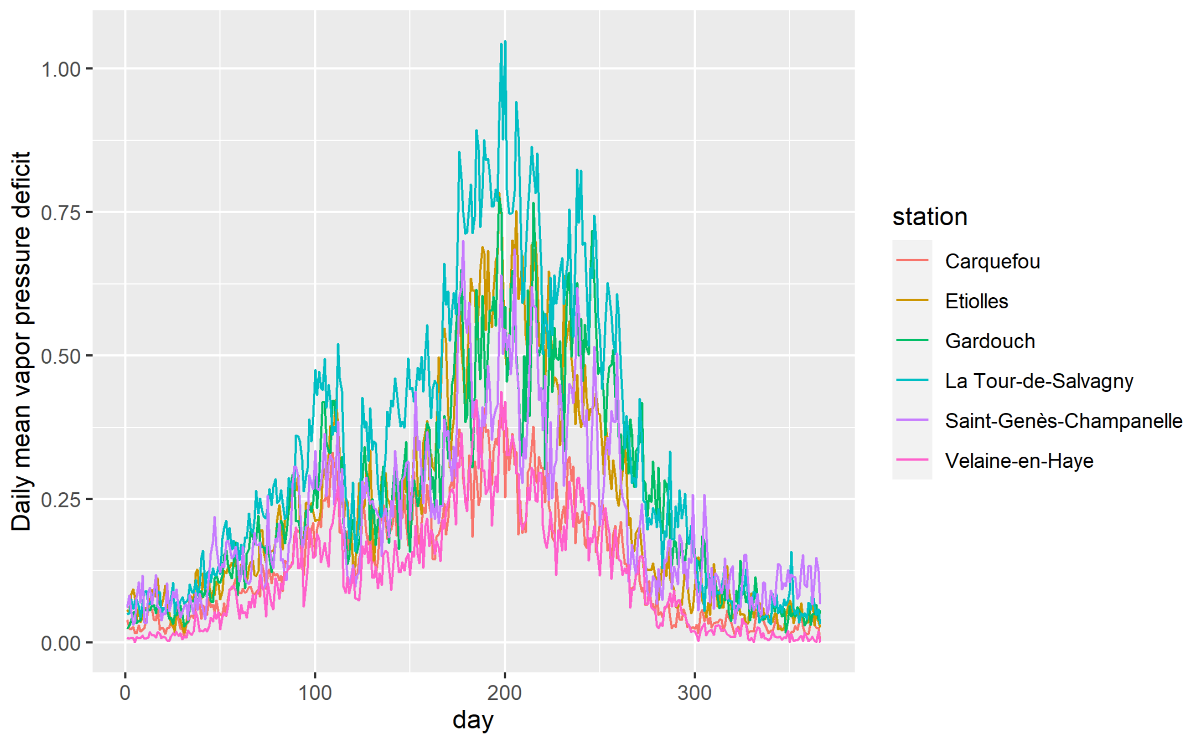


Figure S2: Daily mean observed vapour pressure deficit for the different locations.

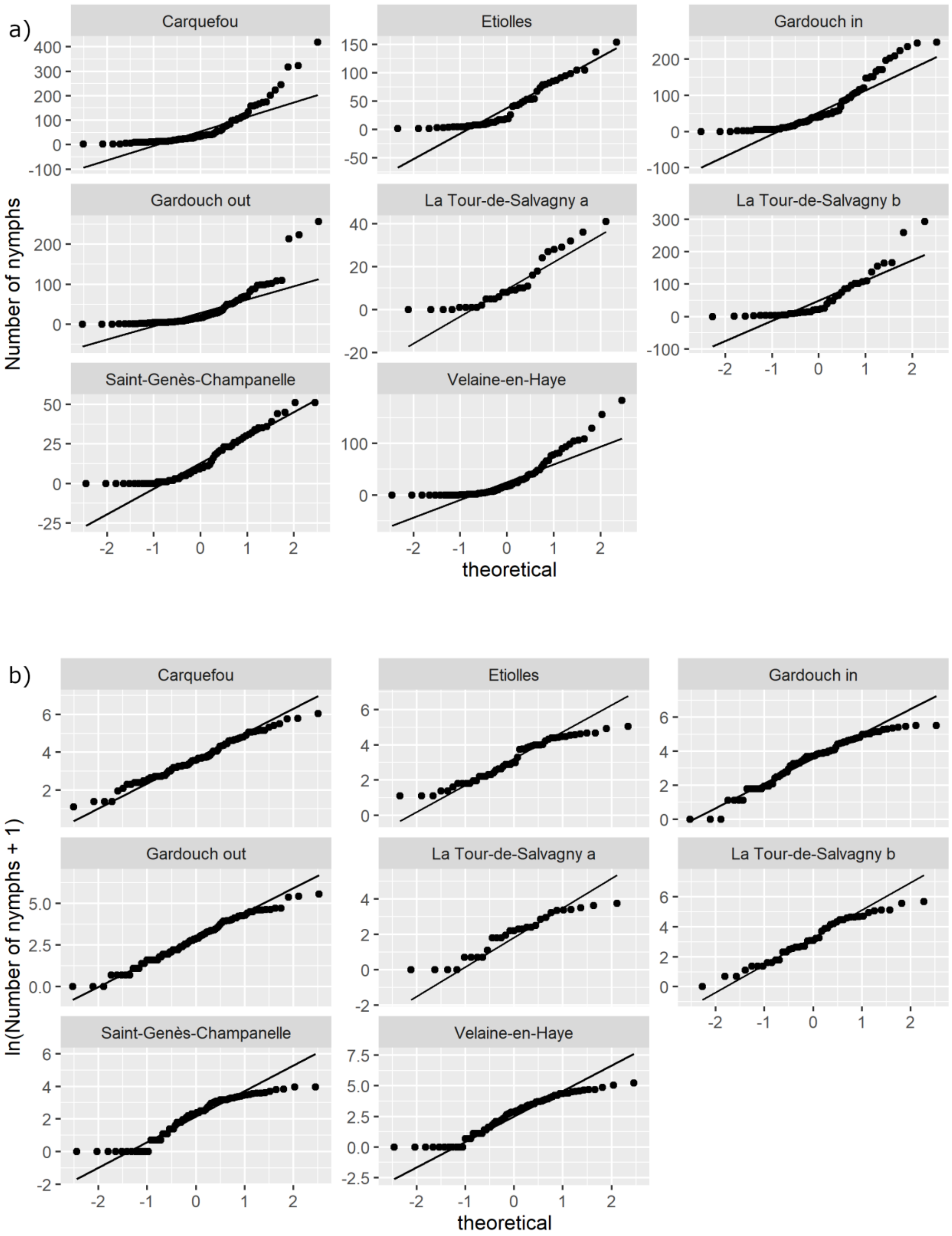


Figure S3: QQ plots of the data for the different sites without (a) or with (b) logarithmic transform.

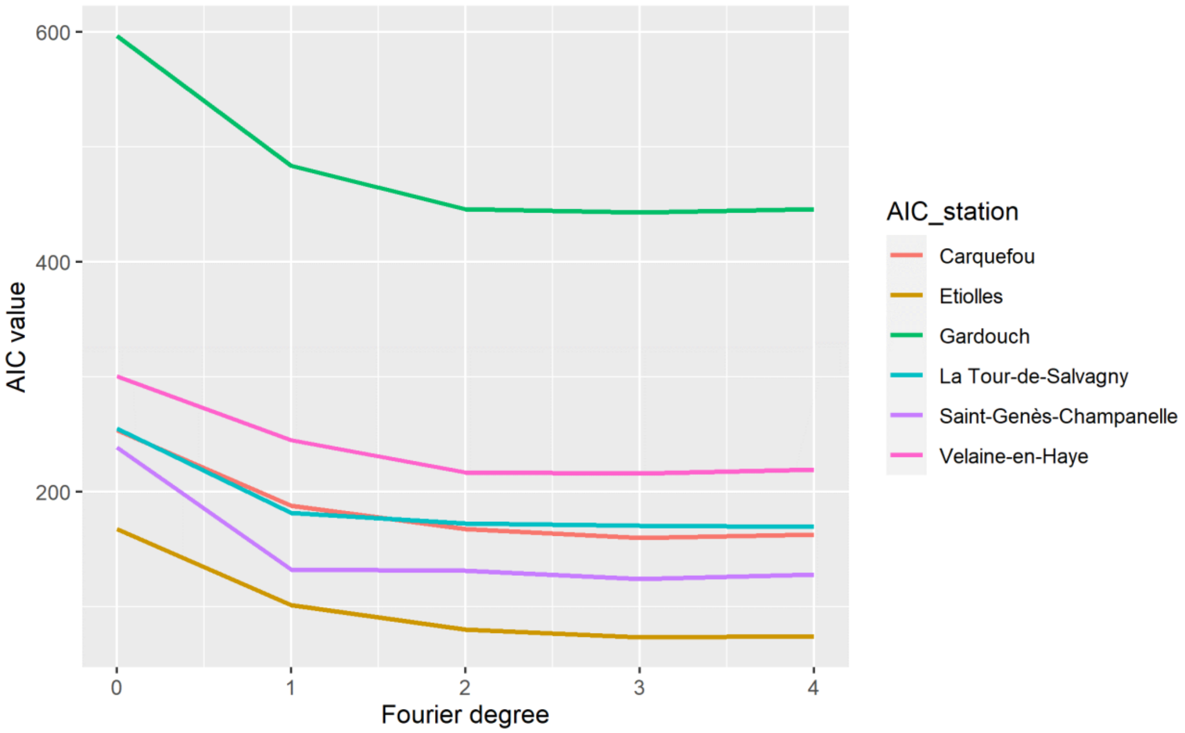


Figure S4: AIC values of harmonic regressions as a function of Fourier degree (parameter *K*) for the different sites. A unique AIC value was computed for La Tour-de-Salvagny and Gardouch sites.

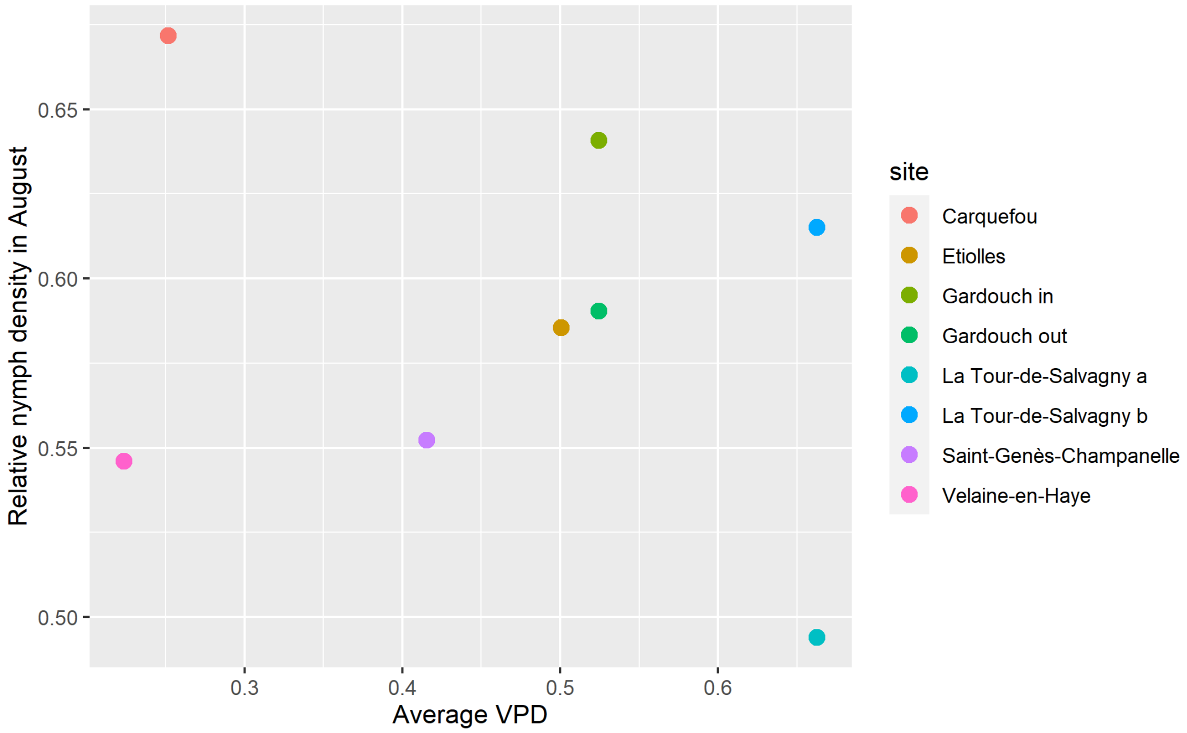


Figure S5: Relationship between Relative nymph density in August and corresponding average vapour pressure deficit.

R script

````
---
title: Modelling Ixodes ricinus phenology
output:
  html_document:
    toc: yes
    toc_depth: '2'
    df_print: paged
  html_notebook:
    toc: yes
    toc_depth: 2
---

# Packages

```{r, message = 'FALSE'}
library(tidyverse)
library(ggplot2)
library(ggrepel)
```


# Data

Uploading tick data

```{r}
ticks <- as_tibble(
  read.csv("data/data_for_regression.csv",
           stringsAsFactors = FALSE))
ticks <- ticks[-which(ticks$IDsite %in% c("9","10","11")),]
```

```{r}
ticks <- ticks %>%
  mutate(date = as.Date(date, "%Y-%m-%d"),
         day_year = as.integer(format(date, "%j")),
         day_radian = day_year / 365 * 2 * pi,
         year = (as.integer(format(date, "%Y"))),
         log_nymph = log(NbNymphe + 1))
```

Corresponding sampling sites and meteorological stations

```{r, message = FALSE}
ls_site_station <- ticks %>%
  group_by(IDsite) %>%
  summarise(
    IDsite = unique(IDsite)) %>%
````
````

```
  mutate(id = row_number(),
         IDstation = c(1,1,2,4,5,7,7,6),
         IDstation_1 = c("S1", "S1", "S2", "S4", "S5", "S6S7", "S6S7", "S8"),
         site = c("La Tour-de-Salvagny a", "La Tour-de-Salvagny b", "Saint-Genès-
Champanelle", "Etiolles", "Carquefou", "Gardouch in", "Gardouch out", "Velaine-en-Haye"),
         station = c("La Tour-de-Salvagny", "La Tour-de-Salvagny", "Saint-Genès-
Champanelle", "Etiolles", "Carquefou", "Gardouch", "Gardouch", "Velaine-en-Haye")) %>%
  select(id, IDsite, IDstation, IDstation_1, site, station)

ticks <- left_join(ticks, ls_site_station)

ls_site_station
```

```{r}
ticks %>%
  group_by(site, station) %>%
  summarise(
    n_collectes = length(unique(date)),
    min_date = min(date),
    max_date = max(date)
  )
```

Distribution of nymph abundance per site after logarithmic transformation

```{r densities log abundances}
ggplot(ticks, aes(x = log(NbNymphe+1))) +
  geom_density() +
  facet_wrap(~site)
ggsave("figures/abundance_log_densities.png")
```

Checking for normality.

```{r qqplot ln abundance}
ggplot(ticks, aes(sample = log(NbNymphe + 1))) +
  stat_qq() +
  stat_qq_line() +
  facet_wrap(~site, scales="free_y") +
  ylab("ln(Number of nymphs + 1)")
ggsave("figures/abundance_log_qqplots.png")
```

Plotting the graph of nymph abundance at each site

````
```{r}
Sys.setlocale("LC_ALL","English")
ticks %>%
  mutate(day_of_year = paste0("2020-", format(date, "%m-%d"))) %>%
  ggplot(., aes(x = as.Date(day_of_year), y = log_nymph, col = factor(year))) +
   geom_point() +
   facet_wrap(~ site) +
   scale_x_date(breaks = function(x) seq.Date(from = as.Date("2020-01-01"),
                                               to = as.Date("2020-12-31"),
                                               by = "1 month"),
                date_labels = "%b") +
  theme(axis.text.x = element_text(angle = 90, hjust = 1)) +
  xlab("") +
  ylab("ln(Number of nymphs + 1)") +
  scale_color_discrete(name = "Year")
ggsave("figures/log_nymph_year_site.png")
```

# Modelling


```{r}
ticks <- ticks %>%
select(IDsite,date,NbNymphe,day_year,day_radian,year,log_nymph,id,IDstation,IDstation_1,
station)
ticks
```

## Choosing degree in Fourier equation by comparing AIC for different models

For Gardouch and La Tour de Salvagny : only the intercept differ between models for two
sites


```{r}
ls_stations <- unique(as.character(ticks$IDstation))

colrs <- c(1,2,3,4,6,7)

AIC_table <- data.frame(
  degre_Fourier=rep(c(0:4),6),
  AIC_station = rep(unique(ticks$station),each=5),
  mdL = NA)

mdL0_1 <- lm(log_nymph ~ 1 + factor(IDsite),
             data = ticks, subset = IDsite %in% c("1a", "1b"))

mdL1_1 <- lm(log_nymph ~  factor(IDsite) +
               cos(day_radian) + sin(day_radian),
````
````

```
           data = ticks, subset = IDsite %in% c("1a", "1b"))

mdL2_1 <- lm(log_nymph ~ factor(IDsite) +
            cos(day_radian) + sin(day_radian) +
            cos(2 * day_radian) + sin(2 * day_radian),
           data = ticks, subset = IDsite %in% c("1a", "1b"))

mdL3_1 <- lm(log_nymph ~ factor(IDsite) +
            cos(day_radian) + sin(day_radian) +
            cos(2 * day_radian) + sin(2 * day_radian) +
            cos(3 * day_radian) + sin(3 * day_radian),
           data = ticks, subset = IDsite %in% c("1a", "1b"))

mdL4_1 <- lm(log_nymph ~ factor(IDsite) +
            cos(day_radian) + sin(day_radian) +
            cos(2 * day_radian) + sin(2 * day_radian) +
            cos(3 * day_radian) + sin(3 * day_radian) +
            cos(4 * day_radian) + sin(4 * day_radian),
           data = ticks, subset = IDsite %in% c("1a", "1b"))

AIC_table$mdL[1] <- AIC(mdL0_1)
AIC_table$mdL[2] <- AIC(mdL1_1)
AIC_table$mdL[3] <- AIC(mdL2_1)
AIC_table$mdL[4] <- AIC(mdL3_1)
AIC_table$mdL[5] <- AIC(mdL4_1)

mdL0_5 <- lm(log_nymph ~ 1 + factor(IDsite),
           data = ticks, subset = IDsite %in% c("6", "7"))

mdL1_5 <- lm(log_nymph ~  factor(IDsite) +
            cos(day_radian) + sin(day_radian),
           data = ticks, subset = IDsite %in% c("6", "7"))

mdL2_5 <- lm(log_nymph ~ factor(IDsite) +
            cos(day_radian) + sin(day_radian) +
            cos(2 * day_radian) + sin(2 * day_radian),
           data = ticks, subset = IDsite %in% c("6", "7"))

mdL3_5 <- lm(log_nymph ~ factor(IDsite) +
            cos(day_radian) + sin(day_radian) +
            cos(2 * day_radian) + sin(2 * day_radian) +
            cos(3 * day_radian) + sin(3 * day_radian),
           data = ticks, subset = IDsite %in% c("6", "7"))

mdL4_5 <- lm(log_nymph ~ factor(IDsite) +
            cos(day_radian) + sin(day_radian) +
            cos(2 * day_radian) + sin(2 * day_radian) +
            cos(3 * day_radian) + sin(3 * day_radian) +
            cos(4 * day_radian) + sin(4 * day_radian),
           data = ticks, subset = IDsite %in% c("6", "7"))
```

```
AIC_table$mdL[21] <- AIC(mdL0_5)
AIC_table$mdL[22] <- AIC(mdL1_5)
AIC_table$mdL[23] <- AIC(mdL2_5)
AIC_table$mdL[24] <- AIC(mdL3_5)
AIC_table$mdL[25] <- AIC(mdL4_5)

pred <- expand.grid(
  date = seq(as.Date("2014-01-01"), as.Date("2019-12-31"), by = "1 day"),
  id = 1:8
)

pred <- full_join(pred, ls_site_station)

pred <- pred %>%
  mutate(day = as.numeric(format(date, "%j")),
         day_radian = day / 365 * 2 * pi)

pred

pred$ln_nNymph <- rep(NA)

pred$ln_nNymph <- rep(NA)


for(i in c(2:4,6)){

 i_station <- ls_stations[i]

mdL0 <- lm(log_nymph ~ 1,
           data = ticks, subset = IDsite == i_station)

mdL1 <- lm(log_nymph ~ cos(day_radian) + sin(day_radian),
           data = ticks, subset = IDsite == i_station)

mdL2 <- lm(log_nymph ~ cos(day_radian) + sin(day_radian) +
             cos(2 * day_radian) + sin(2 * day_radian),
           data = ticks, subset = IDsite == i_station)

mdL3 <- lm(log_nymph ~ cos(day_radian) + sin(day_radian) +
             cos(2 * day_radian) + sin(2 * day_radian) +
             cos(3 * day_radian) + sin(3 * day_radian),
           data = ticks, subset = IDsite == i_station)

mdL4 <- lm(log_nymph ~ cos(day_radian) + sin(day_radian) +
             cos(2 * day_radian) + sin(2 * day_radian) +
             cos(3 * day_radian) + sin(3 * day_radian) +
             cos(4 * day_radian) + sin(4 * day_radian),
           data = ticks, subset = IDsite == i_station)
```

````
AIC_table$mdL[(i - 1)*5+1] <- AIC(mdL0)
AIC_table$mdL[(i - 1)*5+2] <- AIC(mdL1)
AIC_table$mdL[(i - 1)*5+3] <- AIC(mdL2)
AIC_table$mdL[(i - 1)*5+4] <- AIC(mdL3)
AIC_table$mdL[(i - 1)*5+5] <- AIC(mdL4)

}


print(AIC_table)
```


```{r}
AIC_table %>%
  ggplot(., aes(x = degre_Fourier, y = mdL, colour=AIC_station)) +
  geom_line(size = 1) +
  ylab("AIC value") +
  xlab("Fourier degree") +
  scale_color_manual(values=c("#F8766D", "#CD9600", "#00BE67", "#00BFC4",
"#C77CFF", "#FF61CC"))
ggsave("figures/AIC_degree.png")
```

### Best models

```{r}
rm(list = ls()[grep("mdL", ls())])
rm(list = ls()[grep("pred", ls())])
```


```{r, message = FALSE}
pred <- expand.grid(
  date = seq(as.Date("2014-01-01"), as.Date("2021-12-31"), by = "1 day"),
  id = 1:8
)

pred <- full_join(pred, ls_site_station)

pred <- pred %>%
  mutate(day = as.numeric(format(date, "%j")),
         day_radian = day / 365 * 2 * pi)

pred

pred$ln_nNymph <- rep(NA)
```
````
````

````
```{r}
ls_sites <- unique(ticks$IDsite)

mdL_S1 <- lm(log_nymph ~ factor(IDsite) +
               cos(day_radian) + sin(day_radian) +
               cos(2 * day_radian) + sin(2 * day_radian),
             data = ticks, subset = IDsite %in% c("1a", "1b"))

qqnorm(resid(mdL_S1), main = paste("1a", "1b"))
abline(a = 0, b = 1)

pred$ln_nNymph[pred$IDsite %in% c("1a", "1b")] <- predict(mdL_S1, pred[pred$IDsite
%in% c("1a", "1b"),])

mdL_S67 <- lm(log_nymph ~ factor(IDsite) +
               cos(day_radian) + sin(day_radian) +
               cos(2 * day_radian) + sin(2 * day_radian),
             data = ticks, subset = IDsite %in% c("6", "7"))

qqnorm(resid(mdL_S67), main = paste0("6 - 7"))
abline(a = 0, b = 1)

pred$ln_nNymph[pred$IDsite %in% c("6", "7")] <- predict(mdL_S67, pred[pred$IDsite
%in% c("6", "7"),])


for(i in c(3:5, 8)){

i_site <- ls_sites[i]

nm <- paste0("mdL_S", i_site)

mdL <- lm(log_nymph ~ cos(day_radian) + sin(day_radian) +
             cos(2 * day_radian) + sin(2 * day_radian),
           data = ticks, subset = IDsite == i_site)

assign(nm, mdL)

pred$ln_nNymph[pred$IDsite == i_site] <- predict(mdL, pred[pred$IDsite == i_site,])


qqnorm(resid(mdL), main = ls_sites[i])
abline(a = 0, b = 1)

}

pred$ln_nNymph[pred$ln_nNymph < 0] <- 0
```
````

Computation of peak date, minimum date and inflexion point

```{r}
curvs <- pred %>%
  filter(date > "2018-12-31" & date < "2020-01-01") %>%
  group_by(IDsite) %>%
  mutate(dif_1 = ln_nNymph - lag(ln_nNymph, default = NA),
         dif_2 = dif_1 - lag(dif_1, default = NA ),
         dif_1_sign = sign(dif_1) - lag(sign(dif_1), default = NA ),
         dif_2_sign = sign(dif_2) - lag(sign(dif_2), default = NA )
         )

dat_pic <- curvs %>%
  filter(dif_1_sign == -2) %>%
  select( date, id, IDsite, IDstation, site, station,day, ln_nNymph) %>%
  mutate(type_event = "max")

dat_min <- curvs %>%
  filter(dif_1_sign == 2) %>%
  select( date, id, IDsite, IDstation, site, station,day, ln_nNymph) %>%
  mutate(type_event = "min")

dat_inflx <- curvs %>%
  filter(dif_2_sign == -2) %>%
  select( date, id, IDsite, IDstation, site, station,day, ln_nNymph) %>%
  mutate(type_event = "inflx")

curvs_events <- bind_rows(dat_pic, dat_min, dat_inflx) %>%
  arrange(IDsite)

rm(list = c("dat_pic", "dat_min", "dat_inflx"))

curvs_events
```

Creating "events" variable

```{r}
events <- curvs_events %>%
  filter(type_event == "max" & date < "2019-10-01")

events <- bind_rows(
           events,
           curvs_events %>%
             filter(site %in% c("Lyon a", "Lyon b", "Sénart", "Nancy") &
                    type_event == "inflx" & date > "2019-09-01"  & date < "2019-11-01"),
           curvs_events %>%
             filter(site %in% c("Carquefou", "Toulouse int", "Toulouse ext") &
                    type_event == "min" & date > "2019-08-01"  & date < "2019-11-01")
           ) %>%
```

~~~~
  arrange(IDsite)

events
```


Graph superposition with peak date

```{r}
Sys.setlocale("LC_ALL","English")
plot_labs <- curvs %>%
  group_by(site) %>%
  summarise(date = max(date),
            day = max(day),
            ln_nNymph = last(ln_nNymph))

ggplot(curvs, aes(x = date, y = ln_nNymph, colour = site)) +
  geom_line(size = 1) +
  geom_text_repel(data = plot_labs,
                  aes(x = date, y = ln_nNymph, label = site),
                  nudge_x = 30,
                  segment.linetype = 5) +
  geom_point(data = events[events$type_event == "max",],
                  aes(x = date, y = ln_nNymph, colour=site),
             size = 3) +
 scale_x_date(breaks = function(x) seq.Date(from = as.Date("2019-01-01"),
                                            to = as.Date("2019-12-31"),
                                            by = "1 month"),
               date_labels = "%b") +
  ylab("ln(Number of nymphs + 1)") +
  xlab("") +
  theme(axis.text.x = element_text(angle = 90, hjust = 1)) +
  theme(legend.position = "none")
  # scale_color_manual(values=c("#F8766D", "#CD9600", "#7CAE00","#00BE67",
"#00BFC4", "#00A9FF", "#C77CFF", "#FF61CC"))

ggsave("figures/superimposed_abundances.png")
```






```{r}
pred_tick <- pred %>%
  select(date, id, IDsite, IDstation, site, station, day, day_radian, ln_nNymph)

pred_tick <- left_join(pred_tick,
                  ticks %>%
               select(IDsite, date, log_nymph))
```
~~~~

~~~~
Graph with observations and predictions.

```{r}
ggplot(pred_tick, aes(x = date, y = ln_nNymph)) +
  geom_line() +
  geom_point(aes(y = log_nymph, col = "red")) +
  facet_wrap(~ site) +
  theme(legend.position = "none") +
  xlab("Time") +
  ylab("ln(number of nymphs + 1)")
ggsave("figures/ln_nymphs_observed_predicted.png")
```

# Meteorological data


```{r}
meteo <- as_tibble(
  read.csv("data/imputedMeteo_New.csv"))

meteo
```


```{r}
meteo <- meteo %>%
  mutate(date = as.Date(Date))

summary(meteo$date)
```


```{r}
summary(meteo)
```

Computing VPD

```{r}
meteo <- meteo %>%
  mutate(year = format(date, "%Y"),
         day = as.numeric(format(date, "%j")),
         day_radian = day / 365 * 2 * pi,
         VPD=0.6108*exp(17.27*TM_m1/(TM_m1+237.3))*(1-UM_m1/100))
meteo <- meteo[-which(meteo$Site %in% c("S9")),]
```

```{r, message = FALSE}
ls_site_station_meteo <-  meteo%>%
~~~~

~~~~
  group_by(Site) %>%
  summarise(
    Site = unique(Site)) %>%
   mutate(station = c("La Tour-de-Salvagny", "Saint-Genès-Champanelle", "Etiolles",
"Carquefou", "Gardouch", "Velaine-en-Haye")) %>%
   select(Site, station)

meteo <- left_join(meteo, ls_site_station_meteo)

ls_site_station_meteo
```


## Computing daily data means for temperature and VPD


```{r}
meteo_day <- meteo %>%
  group_by(Site, day) %>%
  summarise(station = unique(station),
            date = max(date),
            temp_mean = mean(TM_m1),
            U_min = mean(UN_m1),
            U_mean = mean(UM_m1),
            VPD_mean = mean(VPD))

meteo_day <- meteo_day %>%
  filter(!is.na(temp_mean)) %>%
  mutate(
    temp_tmp = ifelse(temp_mean >= 5,temp_mean, 0),
    temp_cumsum = cumsum(temp_mean),
    temp_cumsum1 = cumsum(temp_tmp),
         U_cumsum  = cumsum(U_mean))
```
```{r}
meteo_day
```


```{r}
ggplot(data = meteo_day, aes(x = day, y = temp_mean, col = station)) +
  geom_line()+
  scale_color_manual(values=c("#F8766D", "#CD9600", "#00BE67", "#00BFC4",
"#C77CFF", "#FF61CC"))+
  ylab("Daily mean temperature")
ggsave("figures/temperature_data.png")
```

```{r}
tick_peak <- events %>%
~~~~

````
  filter(type_event == "max") %>%
  ungroup() %>%
  select(day, station) %>%
  group_by(station) %>%
  summarise(day = unique(day), station = unique(station))

meteo_day$New_Date=as.Date(meteo_day$day, origin = "2021-01-01")

tick_peak <- left_join(
  tick_peak,
  meteo_day %>%
    select(day, New_Date, station, Site, temp_cumsum)
  )
```
Cumulative temperature


```{r}

meteo_day %>%
  filter(day < 220 & temp_cumsum < 2000) %>%
  ggplot(., aes(x = New_Date, y = temp_cumsum, col=station)) +
  geom_line(size = 1) +
  scale_color_manual(values=c("#F8766D", "#CD9600", "#00BE67", "#00BFC4",
"#C77CFF", "#FF61CC")) +
  geom_point(tick_peak,
             mapping = aes(x = New_Date, y = temp_cumsum), size=3) +
    geom_text_repel(data = tick_peak,
                  aes(x = New_Date, y = temp_cumsum, label = station),
                  force = 50,
                  segment.linetype = 5) +
  ylim(0, 2000) +
  xlab("") +
  ylab("Cumulative temperature (°C.d)") +
  theme(legend.position = "none")
ggsave("figures/cumulative_temp_peak.png")

```



```{r}
ggplot(data = meteo_day, aes(x = day, y = VPD_mean, col = station)) +
  geom_line() +
  scale_color_manual(values=c("#F8766D", "#CD9600", "#00BE67", "#00BFC4",
"#C77CFF", "#FF61CC"))+
  ylab("Daily mean vapor pressure deficit")
ggsave("figures/VPD_data.png")

```
````
````

#### Peak date = f(cumulative temperature)

```{r}

date_peak <- events %>%
     filter(type_event == "max") %>%
     ungroup() %>%
     select(day, station) %>%
     group_by(station)
# %>%
#   summarise(day, station)

date_peak <- left_join(
  date_peak,
  meteo_day %>%
    select(day, New_Date, station, Site, temp_mean, temp_cumsum)
  )

date_peak$station

print("Date du pic : ")
date_peak$day

print("Cumul des températures à cette date : ")
date_peak$temp_cumsum

print("Température moyenne à cette date : ")
date_peak$temp_mean
```

#### Mean tick abundance in January = f(mean temperature in January)

```{r}
hiv <- tibble(
  site = unique(ls_site_station$site),
  ln_nNymph_MoyHiv = rep(NA),
  ln_nNymph_Max = rep(NA),
  ln_nNymph_Mean = rep(NA),
  ln_nNymph_MoyHiv_sur_Max = rep(NA),
  ln_nNymph_MoyHiv_sur_Mean = rep(NA),
  teta_MoyHiv = rep(NA))
  # predHiv$NymphHiv <- rep(NA)

for(i in 1:nrow(ls_site_station)){
```

````
  ## Moyenne du ln(nombre de nymphes) pour janvier 2019 pour IDsite i
  hiv$ln_nNymph_MoyHiv[i] <- mean(pred$ln_nNymph[pred$id == i & pred$date > "2020-12-31" & pred$date < "2021-02-01"])
  ## Max du ln(nombre de nymphes) pour toute l'année pour IDsite i
  hiv$ln_nNymph_Max[i] <- max(pred$ln_nNymph[pred$id == i])
  ## Moyenne du ln(nombre de nymphes) pour toute l'année pour IDsite i
  hiv$ln_nNymph_Mean[i] <- mean(pred$ln_nNymph[pred$id == i])
  ## Moyenne du ln(nombre de nymphes) pour janvier divisé par le max pour toute l'année pour IDsite i
  hiv$ln_nNymph_MoyHiv_sur_Max[i] <- hiv$ln_nNymph_MoyHiv[i]/hiv$ln_nNymph_Max[i]
  ## Moy / max / Moy ?
  hiv$ln_nNymph_MoyHiv_sur_Mean[i] <- hiv$ln_nNymph_MoyHiv[i]/hiv$ln_nNymph_Mean[i]
  hiv$teta_MoyHiv[i] <- mean(meteo_day$temp_mean[meteo_day$station == ls_site_station$station[i]  & meteo_day$date > "2020-12-31" & meteo_day$date < "2021-02-01"])

}

model <- lm(hiv$ln_nNymph_MoyHiv_sur_Max ~ hiv$teta_MoyHiv)
param <- coef(model)




hiv
```

```{r}
ggplot(hiv, aes(x = teta_MoyHiv, y = ln_nNymph_MoyHiv_sur_Max, col = site)) +
  xlab("Average temperature (°C)") +
  ylab("Relative nymph density") +
  geom_point(size=3) +
  geom_abline(intercept = -0.206, slope = 0.124, size=1)
ggsave("figures/winter_density_temperature.png")
```

Mean tick in August = f(VPD)

```{r}
aug <- tibble(
  site = unique(ls_site_station$site),
  ln_nNymph_MoyAug = rep(NA),
  ln_nNymph_Max = rep(NA),
  ln_nNymph_MoyAug_sur_Max = rep(NA),
  VPD_MoyAug = rep(NA))

for(i in 1:nrow(ls_site_station)){
````

````
  ## Moyenne du ln(nombre de nymphes) pour aout pour IDsite i
  aug$ln_nNymph_MoyAug[i] <- mean(pred$ln_nNymph[pred$id == i & pred$date > "2020-07-31" & pred$date < "2021-09-01"])
  ## Max du ln(nombre de nymphes) pour toute l'année pour IDsite i
  aug$ln_nNymph_Max[i] <- max(pred$ln_nNymph[pred$id == i])
  ## Moyenne du ln(nombre de nymphes) pour aout divisé par le max pour toute l'année pour IDsite i
  aug$ln_nNymph_MoyAug_sur_Max[i] <- aug$ln_nNymph_MoyAug[i]/aug$ln_nNymph_Max[i]
  aug$VPD_MoyAug[i] <- mean(meteo_day$VPD_mean[meteo_day$station == ls_site_station$station[i]  & meteo_day$date > "2020-07-31" & meteo_day$date < "2020-09-01"])

}

cor(aug$ln_nNymph_MoyAug_sur_Max,aug$VPD_MoyAug)
cor.test(aug$ln_nNymph_MoyAug_sur_Max,aug$VPD_MoyAug)

aug
```

```{r}
ggplot(aug, aes(x = VPD_MoyAug, y = ln_nNymph_MoyAug_sur_Max, col = site)) +
  xlab("Average VPD") +
  ylab("Relative nymph density in August") +
  geom_point(size=3)
ggsave("figures/august_density_VPD.png")
```
````
````
